# Non-HLA antibodies worsen the histological phenotype and prognosis of antibody mediated rejection in kidney allografts

**DOI:** 10.64898/2026.01.09.698585

**Authors:** Emilie Lebraud, Olivier Aubert, Laïla Aouni, Virginia Garcia, Lise Morin, Maëva Eloudzeri, Baptiste Lamarthée, Zulien Zuber, Corentin Ramaugé Parra, Mélanie Try, Maarten Coemans, Adel Abderrahmane, Jean-Luc Taupin, Chloé Jager, Quentin Holleville, Corinne Normand, Frank Bienaimé, Nicolas Garcelon, Fabiola Terzi, Marion Rabant, Dany Anglicheau

**Author notes:** **Corresponding authors:** Dany Anglicheau, Department of Nephrology and Kidney Transplantation, Hôpital Necker., 149, rue de Sèvres, 75015 Paris, France., Emilie Lebraud, Université Paris Cité, Necker-Enfants Malades Institute, INSERM U1151, CNRS UMR8253., 156-160 rue de vaugirard, Paris, France.

## Abstract

**Introduction:** Antibody-mediated rejection (AMR) remains a leading cause of kidney allograft failure, with both HLA and non-HLA antibodies implicated in its pathogenesis. The contribution of non-HLA antibodies (non-HLA Abs) to microvascular inflammation (MVI) and graft outcome, particularly in cases lacking donor-specific anti-HLA antibodies (HLA-DSAs), remains incompletely understood.

**Methods:** We analyzed 571 post-transplant serum samples from 326 patients with histological features of AMR (AMRh) and 164 stable controls. Non-HLA Abs were detected using the previously developed Non-HLA Antibody Detection Immunoassay (NHADIA), and associations were examined with histological lesions, AMRh persistence, and graft outcomes. Biopsies were scored according to Banff 2022 criteria, and patients were stratified by HLA-DSA and NHADIA status.

**Results:** NHADIA values were significantly higher in AMRh patients compared to controls (P=0.0001), regardless of HLA-DSA status. NHADIA values correlated with the severity of glomerulitis, peritubular capillaritis and global MVI scores. In AMRh patients with HLA-DSAs, non-HLA Abs remained independently associated with MVI severity. Follow-up biopsies revealed persistent AMR lesions in patients with both HLA-DSAs and non-HLA Abs. Allograft survival was lowest in double-positive patients, and NHADIA positivity independently predicted graft loss (HR=2.25, 95% CI: 1.03–4.92, P=0.042). Incorporating NHADIA into the Banff classification reclassified 61% of AMRh cases as “double-positive AMRh,” and identified new subgroups with significant prognostic differences.

**Conclusion:** Post-transplant detection of non-HLA antibodies identifies a distinct subset of AMR with more severe histology and worse graft prognosis, particularly when coexisting with HLA-DSAs. Integrating non-HLA Ab testing into current diagnostic frameworks may refine AMR classification and improve risk stratification.

**TRANSLATIONAL STATEMENT:** This study highlights the clinical relevance of non-HLA antibodies, identified using our innovative endothelial cell-based assay (NHADIA), in kidney transplant recipients. Their presence is associated with more severe antibody-mediated rejection (AMR) and poorer graft outcomes, even in the absence of donor-specific HLA antibodies. Incorporating non-HLA antibody detection into routine post-transplant evaluation may allow clinicians to better identify high-risk patients, including those previously classified as DSA-negative AMR. This expanded immunological profiling refines AMR diagnosis, improves risk stratification, and opens new avenues for personalized immunosuppressive strategies, ultimately enhancing long-term graft survival and patient care.

## INTRODUCTION

After kidney transplantation, the Banff International Classification of kidney allograft rejection categorizes alloimmune injury into two main types: T-cell-mediated rejection (TCMR) and antibody-mediated rejection (AMR)^1^, the latter being associated with poor transplant outcome^2,3^. Until recently it was generally accepted that during AMR, pathogenic allo-antibodies were directed against Human Leukocyte Antigens (HLA). Although histological findings suggestive of AMR—such as microvascular inflammation (MVI)—typically correlate with anti-HLA-mediated injury^4^, recent large cohort studies have shown that 40-60% of patients with histological features of AMR are negative for anti-HLA DSAs at the time of biopsy^5–7^. This suggests, among other mechanisms, the potential role of non-HLA antibodies (Abs), as acknowledged in the 2019 Banff classification^1,8^.

The most recent 2022 update of the Banff classification introduced a new diagnostic category: “microvascular inflammation/injury donor-specific antibodies-negative and C4d negative”^9^, further supporting the hypothesis that MVI can result from alternative mechanisms of endothelial injury, including the involvement of non-HLA Abs. This newly defined phenotype has been associated with a higher risk of disease progression and poorer long-term graft survival compared to non-rejection phenotype^10^.

Over the past decades, numerous non-HLA antigens targeted by these Abs have been proposed^11–14^, but their prevalence in transplant patients varies widely depending on the study cohort and detection methods. Available evidence suggests that non-HLA Abs may contribute to kidney allograft rejection^15–18^, and play a broader role in solid organ transplantation^19^. However, whether these Abs act independently or synergistically with HLA-DSAs remains largely unknown.

We previously developed NHADIA (Non-HLA Antibody Detection ImmunoAssay), an endothelial cell-based assay using the CiGEnCΔHLA human glomerular microvascular endothelial cell line, which lacks HLA expression on its surface. This allows for the specific detection of non-HLA Abs, even in patients with circulating HLA-DSAs. In an earlier study, we assessed non-HLA Abs using NHADIA in pre-transplant serum samples of kidney transplant recipients (KTRs)^20^ and found that higher pretransplant levels of non-HLA Abs were associated with an increased risk of post-transplant AMR. Moreover, NHADIA status and HLA-DSAs were found to be independent yet synergistic predictors of AMR^20^.

In the present study, we analyzed 1,980 kidney transplants performed between 2009 and 2021, with serum samples collected at the time of biopsy. We provided evidence that the presence of non-HLA Abs at the time of AMR was associated with MVI lesions and worsens graft prognosis and survival in synergy with HLA-DSA. The aim of this study was to assess the association of non-HLA Abs with the presence, activity and severity of allograft rejection as well as their impact on kidney allograft survival.

## METHODS

### Patient’s selection

We retrospectively reviewed all consecutive adult patients who received a kidney transplant at Necker Hospital (Paris, France) between January 2009 and October 2021, with available serum samples collected at the time of, or immediately before, biopsy (0 days [IQR 1-13.5]) and available follow-up at our institution. The last Banff classification (2022) was used to identify patients with at least one biopsy showing histological features of antibody-mediated rejection (AMR; referred to as AMRh), either with (HLA-DSA+) or without (HLA-DSA-) detectable HLA-DSAs (**Table 1**). Most of patients had one biopsy showing AMRh (N=259 samples) and 67 patients had between two and four biopsies showing AMRh (N=148 samples). A subset of KTRs with one-year protocol biopsies showing no sign of acute rejection (including TCMR or MVI [MVI=0], and no C4d deposition [C4d=0]) were randomly selected as controls. Ultimately, 571 biopsies from 490 KTRs were included: 407 biopsies from 326 patients with AMRh and 164 biopsies from 164 stable patients without evidence of acute rejection (**Figure 1**). Serum samples were obtained from the local Transplantation Biobank (DC-2009-955, Necker Hospital, Paris, France) and all participating patients provided written informed consent.

**Table 1:** Clinical characteristics of AMRh patients. Clinical characteristics of kidney transplant recipients with histological features of antibody-mediated rejection (AMRh), stratified by HLA-DSA status. HLA-DSA with a mean fluorescence intensity (MFI) >500 were considered positive. AMRh: histological features of antibody-mediated rejection; HLA-DSA: Donor-specific anti-human leucocyte antigen antibodies; ECD: Expanded criteria donors; IVIG: Intravenous immunoglobulin.

**Figure 1:**
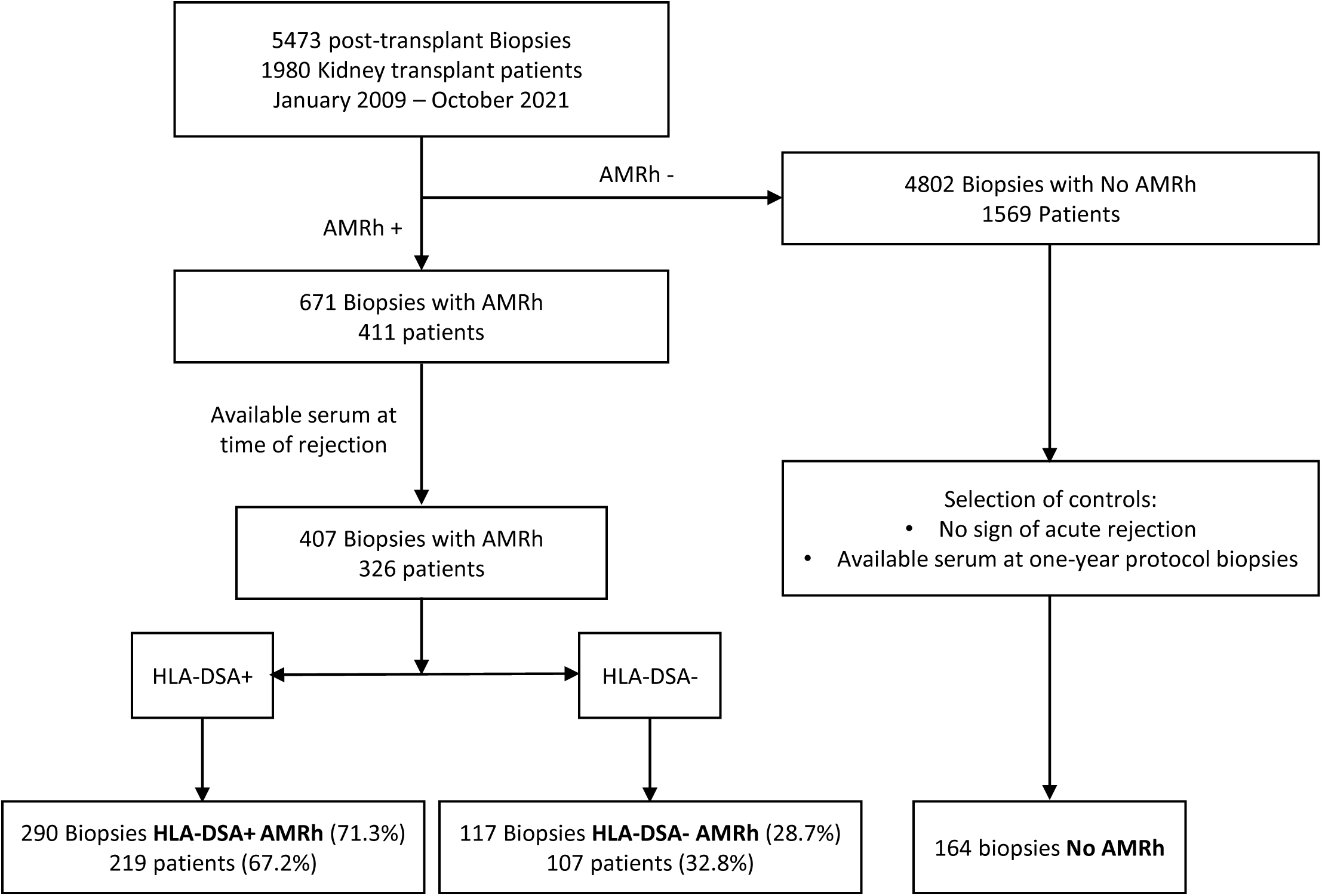
Patient selection and group definition based on AMRh criteria and HLA-DSA status. AMRh: histological features of antibody-mediated rejection; HLA-DSA: Donor-specific anti-human leucocyte antigen antibodies. Biopsies were classified as AMRh if they met one of the following: g + ptc ≥ 2, or g + ptc = 1 with a positive C4d staining, or g + ptc = 0 with TMA or v lesions and a positive C4d staining.

Clinical and biological data for the donors and recipients were retrospectively retrieved from the DIVAT prospective database (Données Informatiques Validées en Transplantation, http://www.divat.fr/), approved by the National French Commission for Bioinformatics Data (CNIL, registration number: 1016618, validated June 8, 2004).

### Histologic assessment and biopsies selection

Biopsy specimens were fixed in formalin, acetic acid and alcohol and embedded in paraffin. Tissue sections were stained with hematoxylin and eosin, Masson’s trichrome, periodic acid–Schiff reagent, and Jones’ stain for light microscopy. C4d immunohistochemical staining was systematically performed using a rabbit anti-human C4d monoclonal Ab (1:200 dilution; Clinisciences).

Protocol biopsies were performed at 3 and 12 months post-transplantation, while indication biopsies were performed for clinical indications. Biopsies were classified using the Banff 2022 update of the Banff classification system^1,9^. The biopsies were deemed inadequate if they contained fewer than 7 glomeruli. Banff elementary lesions were graded from 0 to 3 based on the following histological parameters: peritubular capillaritis (ptc), glomerulitis (g), tubulitis (t), interstitial infiltrate (i), intimal arteritis (v), interstitial fibrosis (ci), tubular atrophy (ct), chronic vascular changes (cv), arteriolar hyalinosis (ah), mesangial matrix expansion (mm), and allograft glomerulopathy (cg)^8^. Microvascular inflammation (MVI) was defined as the sum of g and ptc scores. Banff lesions such as ti, i-IFTA and t-IFTA were not considered, as they were not available for all biopsies.

For the purpose of the study, “AMRh” was defined as biopsies fulfilling histological criteria for AMR, based on Banff scores for g, ptc, v, thrombotic microangiopathy (TMA), cg and C4d deposition^5^. Specifically, biopsies were classified as AMRh if they met one of the following: g + ptc ≥ 2, or g + ptc = 1 with a positive C4d staining, or g + ptc = 0 with TMA or v lesions and a positive C4d staining.

### Detection of HLA-DSA

Details regarding the detection of HLA-DSAs are provided in the **Supplementary Methods**.

### NHADIA

CiGEnCΔHLA cells were obtained and cultured as previously described^20^. Further details regarding the NHADIA experiment are provided in the **Supplementary Methods**. Briefly, NHADIA targets the CiGEnC cells, a conditionally immortalized human microvascular glomerular endothelial cell line, which does not express HLA class I and II molecules following their genetic modification by CRISPR/Cas9 (CiGEnCΔHLA). This cell-based assay is based on the incubation of CiGEnCΔHLA cells with the recipient’s serum. The binding of non-anti HLA antibodies to their targets is detected by flow cytometry, using a secondary anti-human IgG antibody coupled to Alexa Fluor®488.

### Statistical methods

Continuous variables are described as the mean ± standard deviation (SD) or median and IQR. We compared means and proportions between groups by using Mann Whitney test (and Kruskal-Wallis for more than 2 groups), or the Fisher’s exact.

We evaluated the association between the NHADIA levels and each allograft diagnoses. To investigate the association of NHADIA with rejection severity, we compared the mean NHADIA level according to each active and chronic Banff lesion score (ranked from 0 to 3).

Association of pre- and post-transplant factors associated with the levels of NHADIA were assessed using a univariable and multivariable linear regression. Significant factors identified in the univariable analysis were selected for multivariable analysis if the p-value was below 0.20. Parameters included in the final multivariable model were identified using a stepwise backward elimination until each parameter was associated with allograft rejection with a p-value below 0.05.

To identify the optimal prognostic threshold of the NHADIA for graft survival, we used the maximally selected rank statistics method based on the log-rank test. This approach determines the cut-off value that best separates patients into two groups with significantly different risks of allograft loss. The analysis was performed using the maxstat package in R, with the Hothorn–Lausen method used to approximate p-values. Kidney allograft survival was analyzed from the time of transplantation up to a maximum follow-up of 10 years, with graft loss defined as return to dialysis considered the event of interest. For patients who died with a functioning graft, follow-up was censored at the time of death. Survival curves were generated using the Kaplan-Meier method and compared according to NHADIA and DSA status using the log-rank test. Association of NHADIA with kidney allograft adjusted for the presence of HLA-DSAs and histological lesions was performed using a Cox model.

Dendrogram and correlation matrix were built using the ggdendro and Hmisc R packages, respectively. Analyses were performed with R software (version 4.4.1) and GraphPad PRISM® software (version 10.4.2).

## RESULTS

### AMRh and stable patients’ characteristics

According to our inclusion criteria, a total of 571 serum samples were collected at the time of post-transplant biopsies from 326 patients with AMRh and 164 control patients without AMRh (**Figure 1**). Patient and donor characteristics for both AMRh and control groups are summarized in **Table 1** and **Supplementary Table 1**.

Among AMRh patients, those who were HLA-DSA positive (N=219 patients) were significantly younger than HLA-DSA negative AMRh patients (N=107) (P=0.0004) and received kidney allografts from younger donors (P=0.0053). HLA-DSA+ AMRh patients also had a higher frequency of sensitizing events, with a numerically greater proportion having a history of pregnancy, and significantly more previous transplantations (P<0.0001) and blood transfusions (P=0.0066) compared to HLA-DSA-AMRh patients.

Serum creatinine and proteinuria at the time of AMRh diagnosis were comparable between the two groups, as were the proportion of indication versus screening biopsies (**Table 1**).

### Histological characteristics of AMRh and stable patients

We compared the histological characteristics of the 407 biopsies from the AMRh patients according to their HLA-DSA status (**Table 2**). Biopsies from HLA-DSA+ patients exhibited significantly more severe glomerulitis, peritubular capillaritis and MVI compared to those from HLA-DSA-patients (g score=1.4±0.9 vs 1.1±0.8, P=0.0112; ptc score =1.5±1 vs 1.2±1; P=0.003, and MVI=2.9±1.3 vs 2.3±1.3; P=0.0001, respectively). Conversely, interstitial inflammation, intimal arteritis and chronic vascular lesions were more severe in HLA-DSA-biopsies (i score =0.1±0.5 vs 0.3±0.7, P=0.0393; v score =0.1±0.5 vs 0.3±0.3, P=0.0064; and cv score =0.4±1.1 vs 1.7±1, P=0.0045, respectively). Interestingly, the frequency of C4d-positive staining was similar between both groups (P=0.1886), and the C4d Banff scores were also comparable, regardless of HLA-DSA status (P=0.2372) (**Table 2**).

**Table 2:** Histological characteristics of biopsies from AMRh patients. Histological features of biopsies at the time of AMRh diagnosis, stratified by HLA-DSA status. HLA-DSA with a mean fluorescence intensity (MFI) >500 were considered positive. MVI: Microvascular inflammation; AMRh: histological features of antibody-mediated rejection; HLA-DSA: Donor-specific anti-human leucocyte antigen antibodies; TMA: Thrombotic microangiopathy.

The histological characteristics of control group biopsies are presented in **Supplementary Table 2**. In accordance with our inclusion criteria, none of the 164 control biopsies showed MVI lesions or C4d deposition. Most control biopsies exhibited no inflammatory lesions, with the exception of three that showed Borderline changes.

### Post-transplantation non-HLA Abs are more abundant in AMRh patients

The NHADIA assay was performed using the 571 post-transplant serum samples collected from both AMRh and control patients (**Figure 2a**) with CiGEnCΔHLA cells as the targets (**Figure 2b**). The mean NHADIA value was 0.587±0.307, ranging from 0.135 and 3.492, showing large inter-sample variability across the whole population (**Figure 2c**). Univariate and multivariable linear regression analyses identified prior transplantation (ß=0.0723; 95%CI=0.0127-0.1320; P=0.0176) and recipient age (ß=0.0019; 95%CI=0.0002-0.0037; P=0.0279) as the only independent determinants of the NHADIA value (**Supplementary Figure 1**).

**Figure 2:**
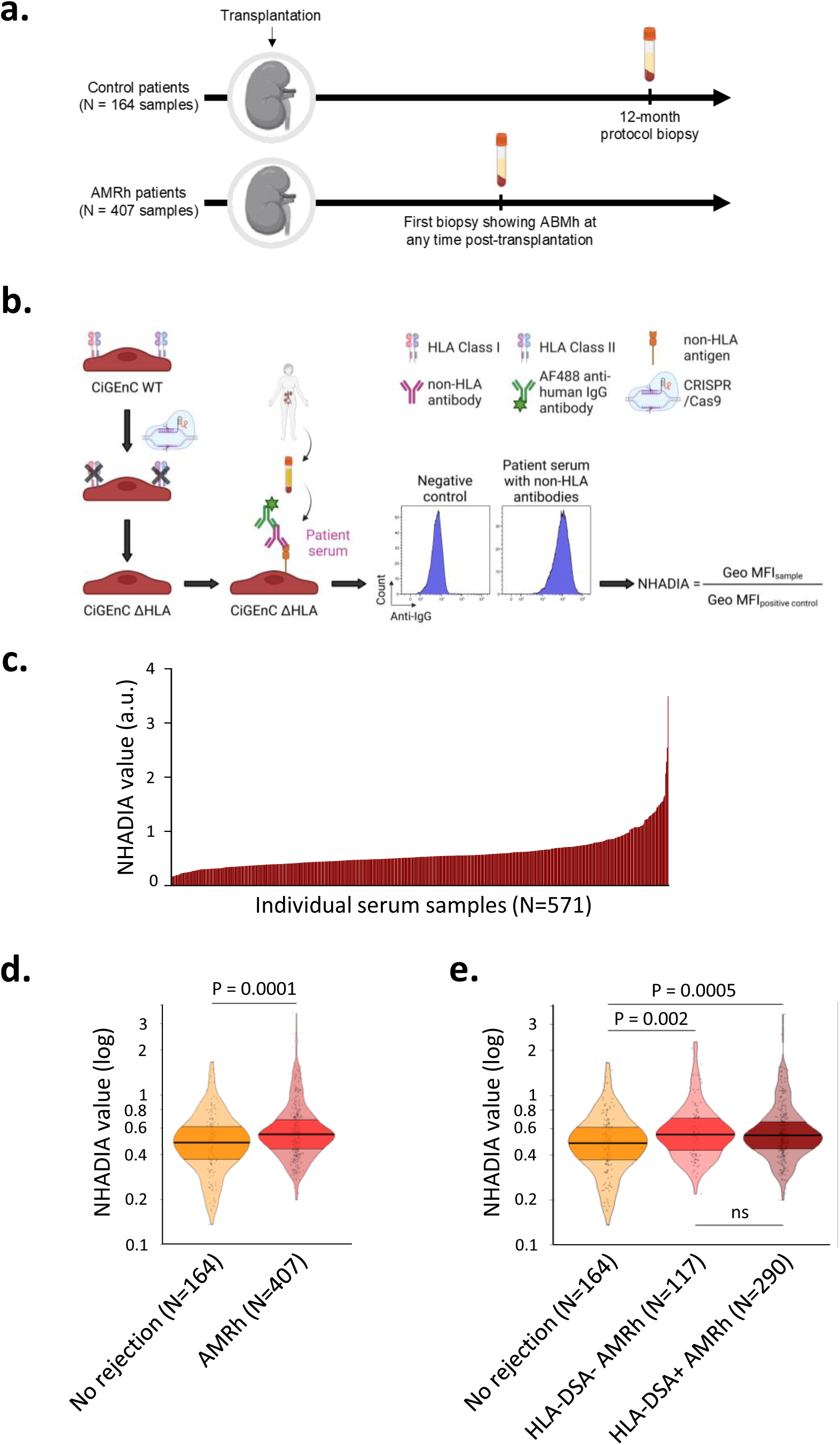
Association of non-HLA antibodies assessed by NHADIA with rejection according to HLA-DSA status. **a**. Study design for control (N=164) and AMRh (N=326) patients. Mo: Months **b**. Overview of the Non-HLA Antibody Detection Immunoassay (NHADIA) applied to 571 post-transplant serum samples. **c**. NHADIA value distribution (a.u.) among control and AMRh patients. **d**. NHADIA value distribution (log scale) in control vs. AMRh patients. **e**. NHADIA values (log scale) in control patients, HLA-DSA-AMRh patients and HLA-DSA+ AMRh patients. AMRh: histological features of antibody-mediated rejection; MFI: Mean fluorescence intensity; HLA-DSA: Donor-specific anti-human leucocyte antigen antibodies; NHADIA: Non-HLA Antibody Detection Immunoassay; a.u.: arbitrary units; P values from Mann-Whitney tests.

NHADIA values were significantly higher in AMRh biopsies compared to controls (P=0.0001) (**Figure 2d**). When AMRh cases were stratified by HLA-DSA status, NHADIA values were significantly higher in both HLA-DSA+ and HLA-DSA-groups compared to controls (P=0.0005 and P=0.002, respectively) with similar NHADIA values between the two AMRh groups (**Figure 2e**). The majority of AMRh patients were positive for both antibody types (HLA-DSA+ NHADIA+, 61%), whereas only 20% of controls showed positivity for either HLA-DSAs or non-HLA Abs (**Supplementary Figure 2**).

### NHADIA values are associated with MVI severity

We evaluated the association between NHADIA values and allograft histology across the full cohort. Unsupervised clustering analysis revealed that NHADIA levels clustered with histological features of MVI (i.e. g and ptc) (**Figure 3a**). A correlation matrix confirmed the significant associations between NHADIA values and glomerulitis (P=0.0001), as well as peritubular capillaritis (P=0.0451) (**Figure 3b**). Higher NHADIA values were observed with more severe lesions of glomerulitis (P=0.0008), peritubular capillaritis (P=0.0111) and MVI (g+ptc, P=0.0012) (**Figure 3c**). Additionally, NHADIA values were associated with tubulitis (P=0.0081) and interstitial inflammation (P=0.0478) and tubulitis + interstitial inflammation (P=0.0218) (**Figure 3d**). No association was observed with chronic histological lesions (**Supplementary Figure 3**).

**Figure 3:**
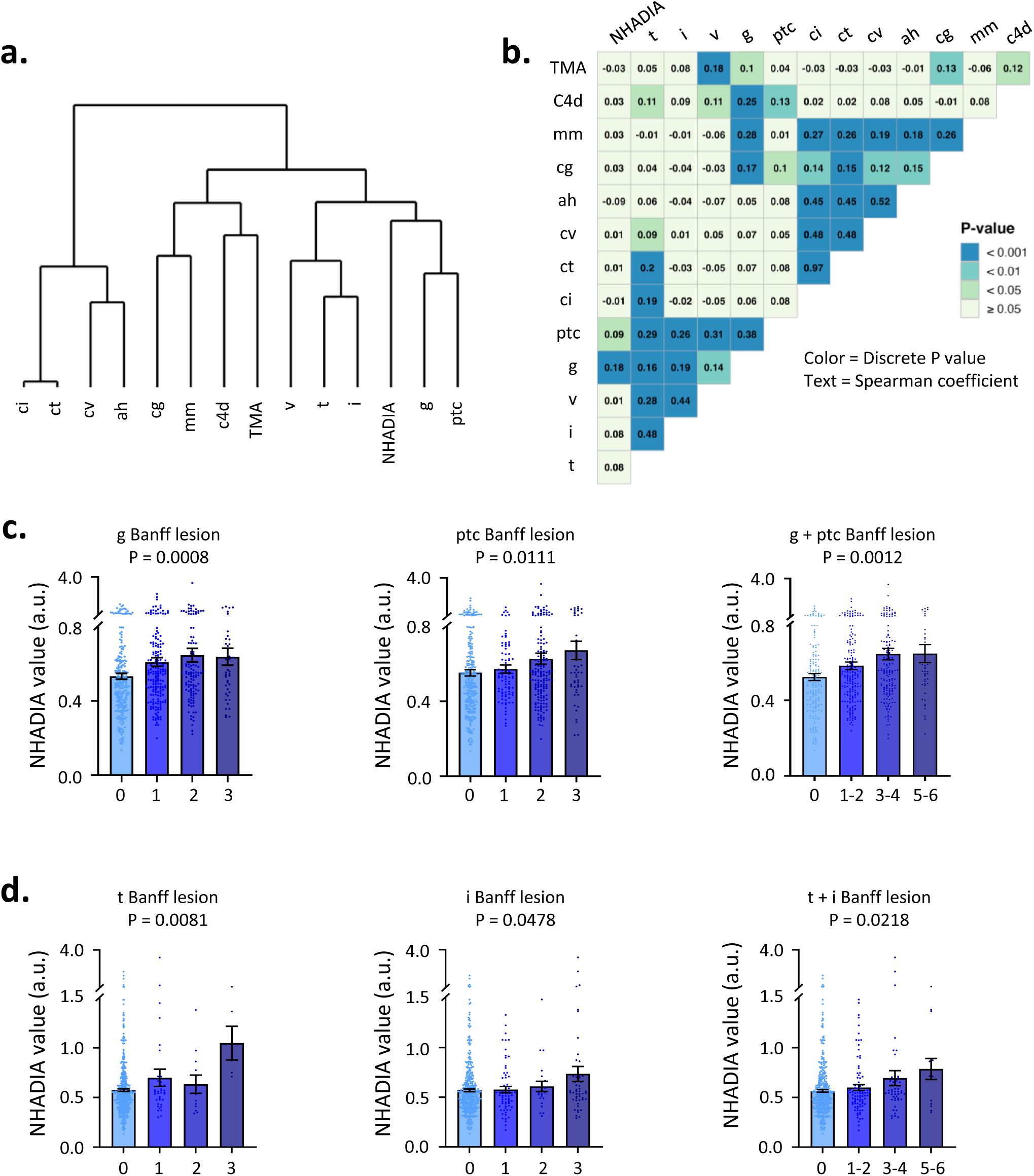
Association of NHADIA with AMR and TCMR features. **a**. Dendrogram of unsupervised hierarchical clustering based on NHADIA quartiles and Banff elementary lesions observed at time of post-transplantation biopsies (control and AMRh patients). The vertical axis of the dendrogram represents the distance or dissimilarity between clusters. **b**. Correlation matrix between NHADIA and Banff elementary lesions (control and AMRh patients). Colors in the squares represent p values and values in the squares represent the Spearman’s Rho. **c**. Mean NHADIA values (a.u.) by g, ptc and g + ptc scores in control and AMRh patients. **d**. Mean NHADIA values (a.u.) by t, i and t + i scores in control and AMRh patients. NHADIA: Non-HLA Antibody Detection Immunoassay; a.u.: arbitrary units; AMRh: histological features of antibody-mediated rejection; TMA: Thrombotic microangiopathy; ah: arteriolar hyalinosis; mm: mesangial matrix expansion; cg: allograft glomerulopathy; cv: chronic vascular changes; ct: tubular atrophy; ci: interstitial fibrosis; v: vasculitis; i: interstitial infiltrate; t: tubulitis; ptc: peritubular capillaritis; g: glomerulitis. P values from two-sided Kruskal-Wallis test. Data are presented as mean values ± SEM.

To isolate the impact of non-HLA Abs, we restricted the analysis to AMRh biopsies from HLA-DSA+ patients. NHADIA values were still significantly associated with glomerulitis (P=0.0245), peritubular capillaritis (P=0.0189), and MVI (P=0.0012) (**Figure 4a**), as well as with tubulitis (P=0.0214), interstitial inflammations (P=0.0329) and tubulitis + interstitial inflammation (P=0.0052) (**Figure 4b**).

**Figure 4:**
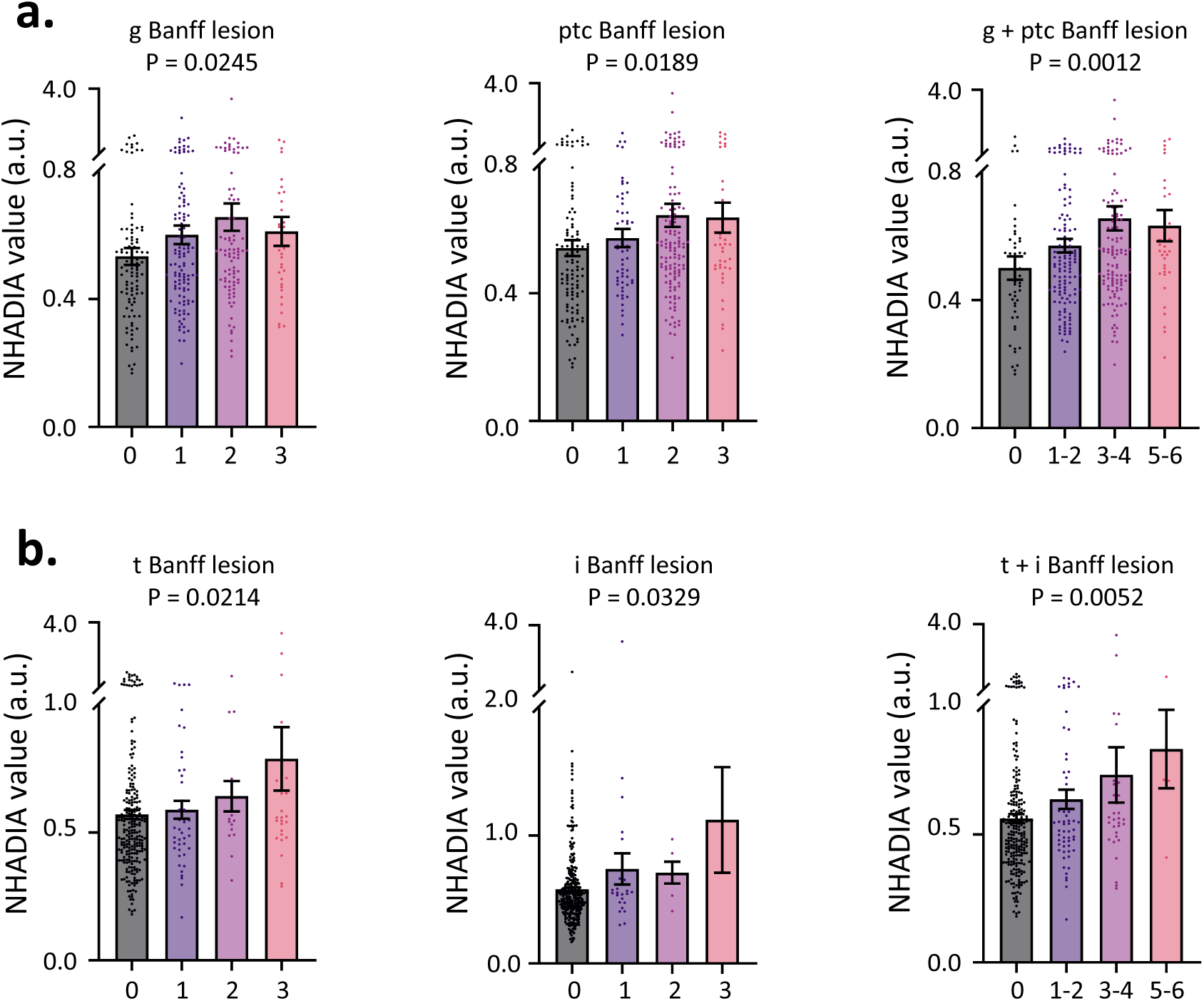
NHADIA association with AMR and TCMR lesions in HLA-DSA+ patients. **a**. Mean NHADIA values (a.u.) by g, ptc and g + ptc scores in HLA-DSA+ control and AMRh patients. **b**. Mean NHADIA values (a.u.) by t, i and t + i scores in HLA-DSA+ control and AMRh patients. AMRh: histological features of antibody-mediated rejection; g: glomerulitis; ptc: peritubular capillaritis; t: tubulitis; i: interstitial infiltrate. P values from two-sided Kruskal-Wallis test. Data are presented as mean values ± SEM.

### Non-HLA Abs synergize with HLA-DSAs to influence AMRh resolution and allograft prognosis

Non-HLA Abs detected at the time of AMRh were associated with a more severe AMR histological phenotype, especially in the presence of HLA-DSAs. Using maximally selected rank statistics, we identified an optimal NHADIA threshold of 0.386 to define positivity, based on its association with the rate of graft loss. Among our 326 AMRh patients, we analyzed 157 follow-up biopsies performed at a median of 7.4 [5.3-8.9] months after the index AMRh biopsy (**Figure 5a**). The best histological resolution was observed in patients negative for both HLA-DSAs and non-HLA Abs. Conversely, patients positive for both showed significantly higher g scores (**Figure 5b**), ptc scores (P=0.0011) (**Figure 5c**) and MVI scores (P=0.0112) (**Figure 5d**) in follow-up biopsies, resulting in persistent AMRh diagnosis in 54% of cases (P=0.0444) (**Figure 5e**).

**Figure 5:**
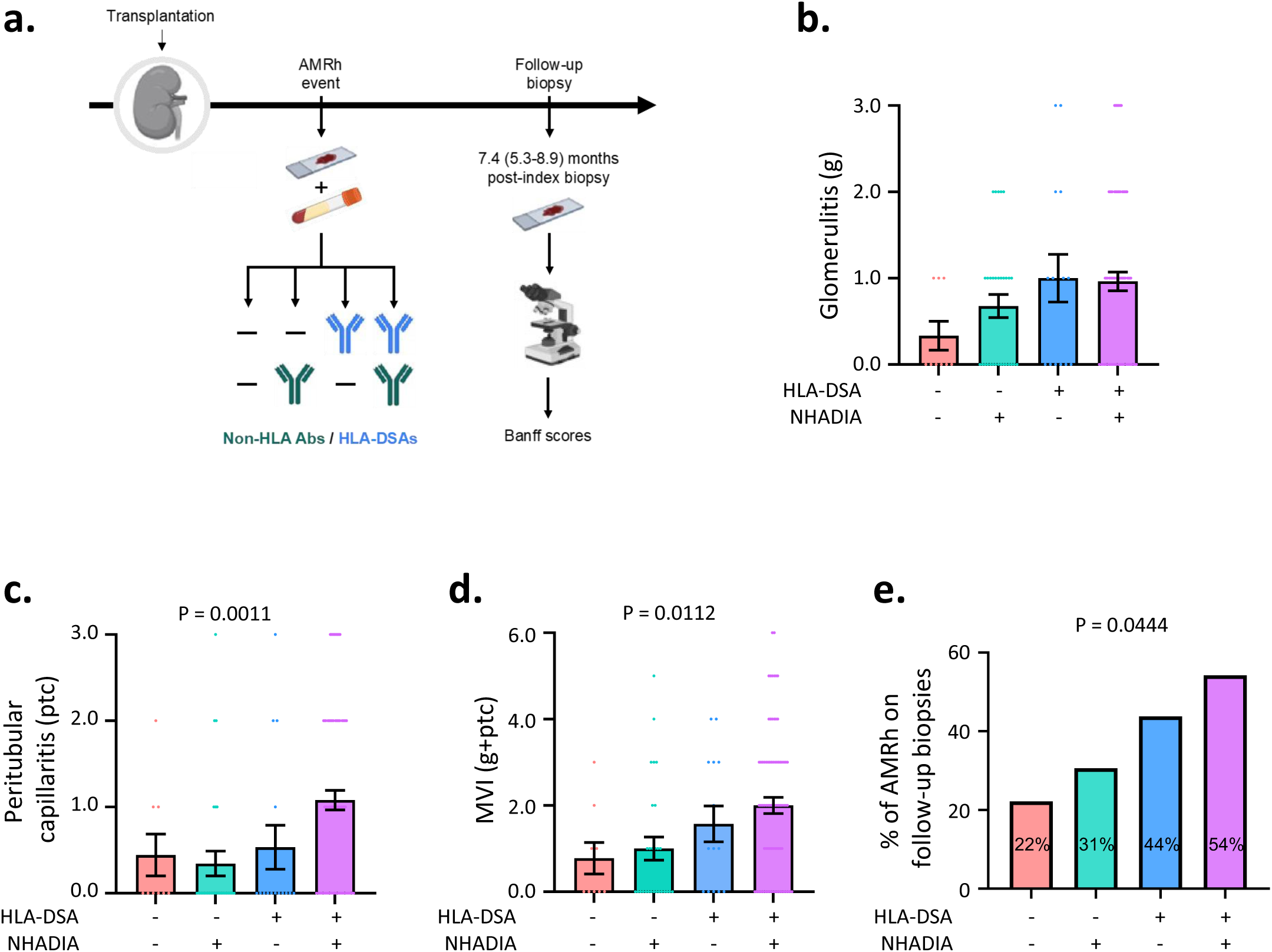
NHADIA and HLA-DSA synergize in AMR persistence. **a**. Design of follow-up biopsy analysis. One hundred and fifty-seven follow-up biopsies performed at a median of 7.4 [5.3-8.9] months after a first AMRh event were analyzed and graded from zero to three according to the Banff histological parameters. Index biopsies were stratified into four groups according to their NHADIA and HLA status. Mean g score (**b**), ptc score (**c**) and total MVI (g+ptc) score (**d**) of follow-up biopsies in the four groups. **e**. Frequency of AMRh persistence among groups. AMRh: histological features of antibody-mediated rejection; MVI: Microvascular inflammation; HLA-DSA: Donor-specific anti-human leucocyte antigen antibodies; g: glomerulitis; ptc: peritubular capillaritis. P values from two-sided Kruskal-Wallis test. Data are presented as mean values ± SEM.

### Non-HLA Abs negatively impact graft survival alongside HLA-DSAs

We next evaluated the association between non-HLA Abs and long-term kidney allograft survival. Patients were stratified into four groups based on HLA-DSA and non-HLA Ab status. Non-HLA Abs were significantly associated with poorer graft survival (P=0.0257) (**Figure 6a**), as were HLA-DSAs (P<0.0001) (**Figure 6b**).

**Figure 6:**
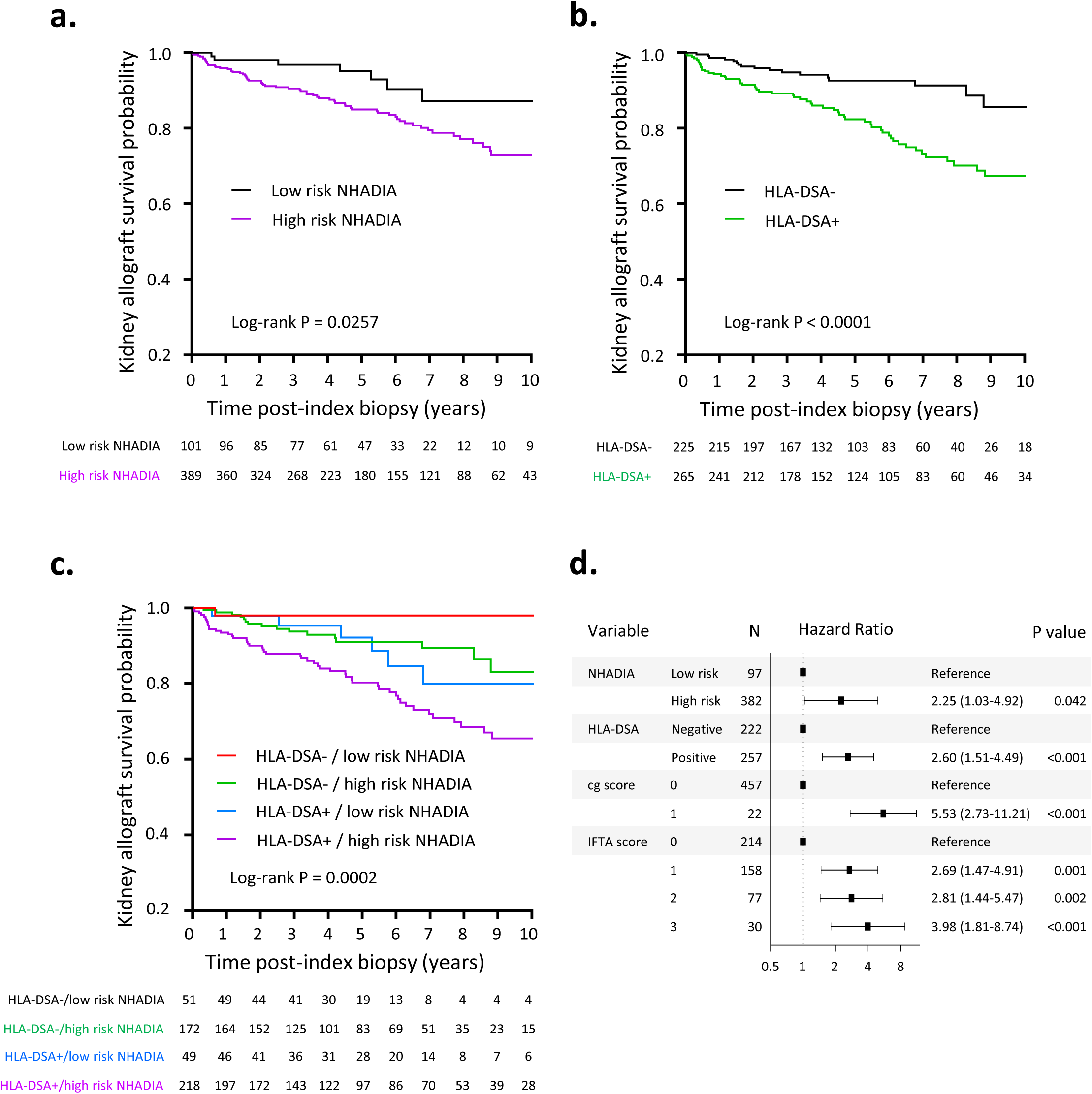
Impact of non-HLA antibodies and HLA-DSA on graft survival. Kaplan-Meier curves by NHADIA (**a**) and HLA-DSA (**b**) status. High risk NHADIA patients were defined by a NHADIA value > 0.3859. P values from log-rank test. **c**. Kaplan-Meier curve among control and AMRh patients stratified by NHADIA and HLA-DSA status. P values from log-rank test. **d**. Multivariable Cox regression of NHADIA adjusted for anti-HLA-DSA and histology. HLA-DSA: Donor-specific anti-human leucocyte antigen antibodies; NHADIA: Non-HLA Antibody Detection Immunoassay; cg: allograft glomerulopathy; IFTA: interstitial fibrosis and tubular atrophy.

Outcomes differed significantly across groups: patients without antibodies had the best survival, those with both had the worst, and intermediate outcomes were seen in patients with only one type of antibody (P=0.0002) (**Figure 6c**). In multivariable Cox regression, NHADIA positivity was independently associated with an increased rate of allograft loss (HR=2.25, 95% CI: 1.03–4.92, P=0.042), adjusting for HLA-DSA status, cg score, and IFTA score (**Figure 6d**).

### Incorporating non-HLA Abs may refine AMR diagnosis and prognosis

To evaluate whether adding NHADIA results to current AMR diagnostic criteria could improve classification and risk stratification, we applied the 2022 Banff classification and incorporated non-HLA Abs status based on the NHADIA threshold (**Figure 7a**). This approach allowed us to define three additional subcategories: AMRh DSA-NHADIA-, AMRh DSA+ NHADIA-, and AMRh DSA-NHADIA+. We also introduced a new category, “Double Positive AMRh”, corresponding to ABMRh cases with both circulating HLA-DSAs and non-HLA Abs. An alluvial diagram (**Figure 7b**) illustrates the reclassification potential: 248 biopsies (61% of total biopsies) originally labeled as AMRh could be reclassified as Double Positive AMRh. Notably, 85% of MVI C4d-DSA-biopsies, could be reclassified as AMRh DSA-NHADIA+ (g+ptc ≥ 2 C4d-DSA-, but NHADIA+).

**Figure 7:**
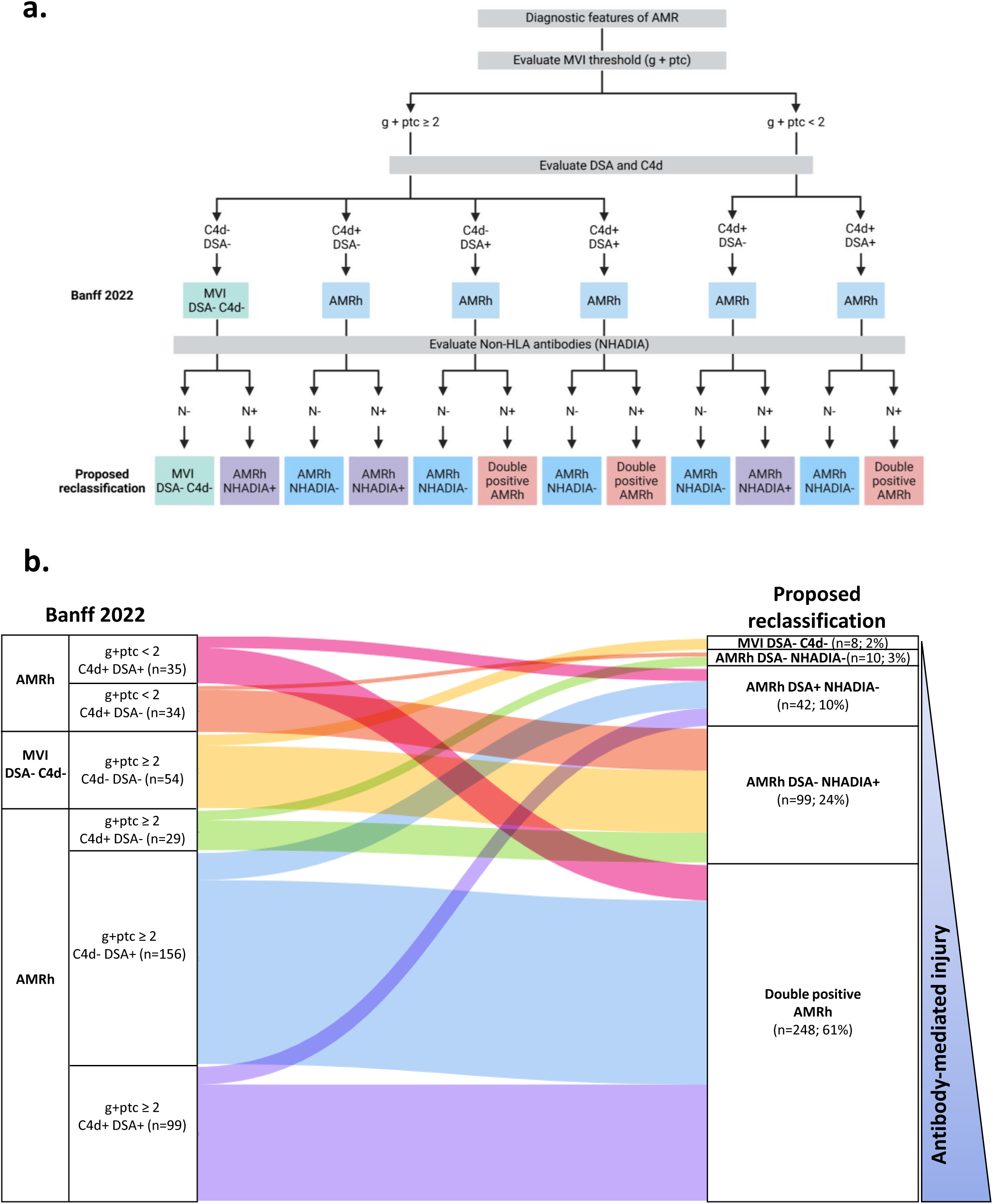
Integration of non-HLA antibodies in AMR classification. **a**. Reclassification of AMRh biopsies using NHADIA, HLA-DSA, and C4d status compared to Banff 2022 classification. N refers to NHADIA **b**. Alluvial diagram showing diagnosis transitions between classifications. MVI: Microvascular inflammation; AMRh: histological features of antibody-mediated rejection; HLA-DSA: Donor-specific anti-human leucocyte antigen antibodies; NHADIA: Non-HLA Antibody Detection Immunoassay; g: glomerulitis; ptc: peritubular capillaritis.

## DISCUSSION

The role of non-HLA Abs is increasingly recognized in AMR, but whether they act independently or synergistically with HLA-DSAs remains unclear. Moreover, the lack of readily available assays limits their identification in clinical practice and prevents the use of antibody-targeting therapies in patients who may benefit from them.

To address these gaps, we previously developed a cellular assay to assess non-HLA antibody burden in AMR patients^20^. Using the CiGEnC human glomerular endothelial cell line as a target, we established an endothelial cell-based assay. To eliminate HLA interference, CiGEnC cells were genetically modified via CRISPR/Cas9 to delete *B2M* and *CIITA*, resulting in CiGEnCΔHLA cells. These cells, phenotypically identical to the parental line, enabled the development of NHADIA (Non-HLA Antibody Detection ImmunoAssay). NHADIA quantifies the burden of non-HLA endothelial antibodies, and we previously demonstrated that pretransplant non-HLA Ab burden assessed by NHADIA predicts the risk of developing AMRh after kidney transplantation^20^.

Here, we provide evidence that post-transplant circulating non-HLA Abs, assessed at the time of biopsy, are associated with MVI in kidney allografts, exacerbate histological lesions in HLA-DSA+ AMRs, and correlate with kidney graft loss independently of HLA-DSAs presence and chronic histological lesion severity.

The causal role of non-HLA Abs in AMR remains debated. It is still largely unknown whether non-HLA Abs represent a nonspecific response to allograft injury without intrinsic pathogenicity or act as active mediators of immune injury^21^. Our results support a pathogenic role for non-HLA Abs detected by NHADIA. First, in terms of temporality, our previous study demonstrated that non-HLA Abs precede the occurrence of MVI; pre-existing non-HLA Abs were associated with post-transplant AMRh independently of HLA-DSAs^20^. Second, in this study, we observed for the first time a dose-dependent effect: the burden of non-HLA Abs correlated with the histological severity of MVI, suggesting a specific, deleterious effect on graft endothelium. Third, consistent with our previous findings on pre-transplant non-HLA Abs^20^, prior exposure to a kidney allograft was the main determinant of post-transplant NHADIA positivity, supporting a role for allo-sensitization to minor histocompatibility antigens—an immune response known to affect long-term graft outcomes^22,23^. Interestingly, unlike anti-HLA Abs, neither pregnancy nor blood transfusion history were found to be associated with the appearance of non-HLA Abs, suggesting that NHADIA-detected Abs may target antigens restricted to kidney tissue. Finally, our study suggests that the non-HLA Ab burden at the time of biopsy exacerbates graft lesions in HLA-DSA+ AMRs and is associated with graft loss independently of HLA-DSAs presence and chronic lesion severity, further supporting a specific pathogenic effect.

Notably, we observed that AMRh was associated with increased non-HLA Abs levels, independently of HLA-DSA status. In other words, even classic HLA-DSA+ AMRs were associated with increased non-HLA Ab burden. This highlights the limitations of oversimplifying the pathophysiological mechanisms of MVI and the importance of considering alternative pathways, including non-HLA Ab–mediated and Ab-independent mechanisms. These findings emphasize the redundant and deeply interconnected nature of the immune system, wherein diverse injurious mechanisms can converge to produce similar histological features.

Our results also reveal that a minority of AMRh patients exhibit neither HLA-DSAs nor non-HLA Abs, supporting the existence of Ab-independent mechanisms of endothelial injury. Several studies have implicated natural killer (NK) cells in MVI lesions via the missing-self mechanism^7,24,25^, or monocyte-mediated graft injury due to SIRPα/CD47 mismatch between donor and recipient^26,27^. These mechanisms are not mutually exclusive, and MVI in allografts likely results from a combination of Ab-dependent and Ab-independent pathways.

Characterizing these mechanisms of allograft injury is critical to defining optimal therapeutic strategies, particularly in a context where the standard of care for AMR remains poorly defined due to the lack of effective treatments. In this regard, incorporating our findings into the Banff classification of AMR could not only expand the recognition of Ab involvement to more patients—including those with MVI but no HLA-DSAs—but also refine prognostication. Specifically, patients with both HLA-DSAs and a high non-HLA Ab burden experience a greater risk of graft loss.

Nonetheless, our study has several limitations. First, NHADIA is based on a genetically modified cell line and may not detect truly donor-specific non-HLA antibodies. However, our data indicate that NHADIA results have both diagnostic and prognostic value in kidney transplant recipients. Second, this study was conducted in a single-center retrospective cohort, lacking external validation in an independent population. Third, the optimal NHADIA cut-off for predicting graft loss was determined using a data-driven method (maximally selected rank statistics) within the same dataset; hence, its prognostic value should be interpreted cautiously until prospectively validated. Finally, although multivariable models adjusted for key histological and immunological covariates, residual confounding inherent to observational designs cannot be entirely excluded.

In summary, our retrospective study of over 490 carefully characterized kidney transplant recipients highlights the relevance of non-HLA antibody burden, as assessed by our novel NHADIA test, as an independent immunological risk factor for graft loss. Identifying patients with high non-HLA antibody burden at the time of rejection not only improves understanding of the underlying mechanisms and informs personalized treatment strategies, but also enhances risk stratification for future graft loss. External validation in larger, more diverse cohorts is now needed to confirm these findings.

## DISCLOSURE STATEMENT

The authors have declared that no conflict of interest exists

## ACKNOWLEDGMENT

The authors thank J. Megret and C. Cordier, who are part of the staff of the cytometry core facility of SFR Necker. Figures 2A, 2B, 5A and 7A were created with BioRender.com.

## DATA AVAILABILITY

All individual data are provided in the manuscript and the supplementary material.

## AUTHOR CONTRIBUTIONS

EL, MR and DA designed the study, created the figures and drafted the paper. EL, LA, JZ, MT, MC, AA, JLT, FB, NG, FT, MR and DA contributed to patient inclusion and data collection. MR reviewed all the kidney biopsies. BL constructed the CiGEnCΔHLA cell line. EL, VG, ME, CJ, QH and CN performed NHADIA assay and EL analyzed flow cytometry results. OA, LM and CRP conducted the statistical analyses. All authors reviewed and approved the final version.

## FIGURE AND TABLE LEGENDS

**Supplementary Table 1**: **Clinical characteristics of AMRh and control patients.** AMRh: histological features of antibody-mediated rejection; HLA-DSA: Donor-specific anti-human leucocyte antigen antibodies; ECD: Expanded criteria donors; IVIG: Intravenous immunoglobulin.

**Supplementary Table 2**: **Histological characteristics of 12-months protocol biopsies from control patients.** MVI: Microvascular inflammation; TMA: Thrombotic microangiopathy.

**Supplementary Figure 1:**
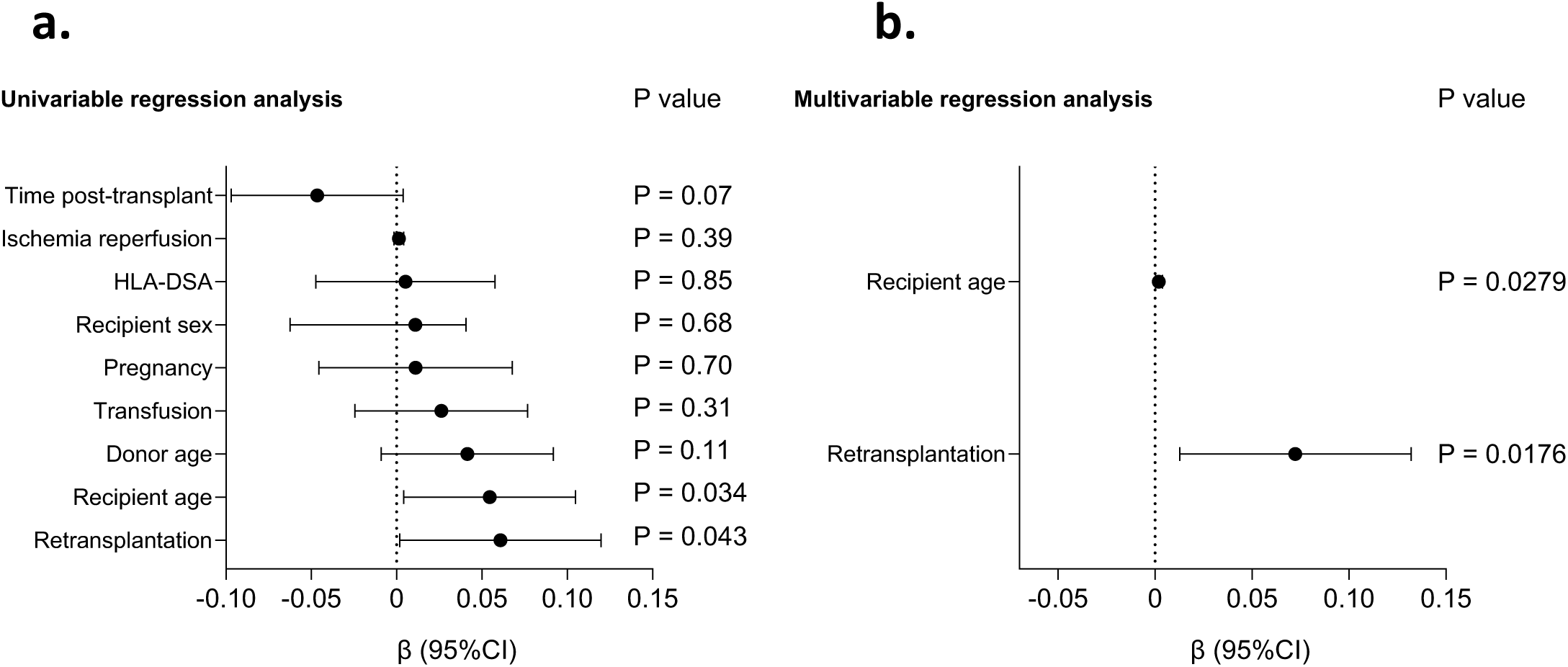
Determinants of NHADIA results at time of rejection. Univariate (**a**) and multivariable (**b**) linear regression analysis of determinants of NHADIA results measured post-transplantation in control and AMRh patients. AMRh: histological features of antibody-mediated rejection; HLA-DSA: Donor-specific anti-human leucocyte antigen antibodies; NHADIA: Non-HLA Antibody Detection Immunoassay.

**Supplementary Figure 2:**
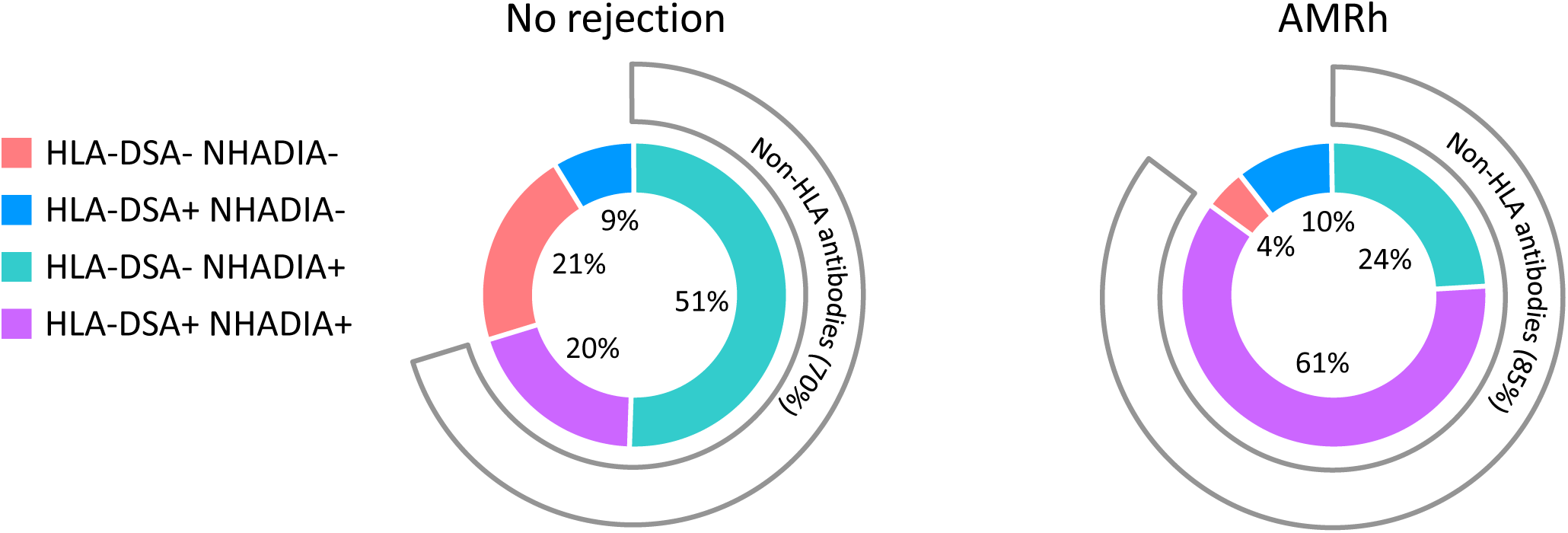
Biopsies distribution by NHADIA and HLA-DSA status. AMRh: histological features of antibody-mediated rejection; HLA-DSA: Donor-specific anti-human leucocyte antigen antibodies; NHADIA: Non-HLA Antibody Detection Immunoassay.

**Supplementary Figure 3:**
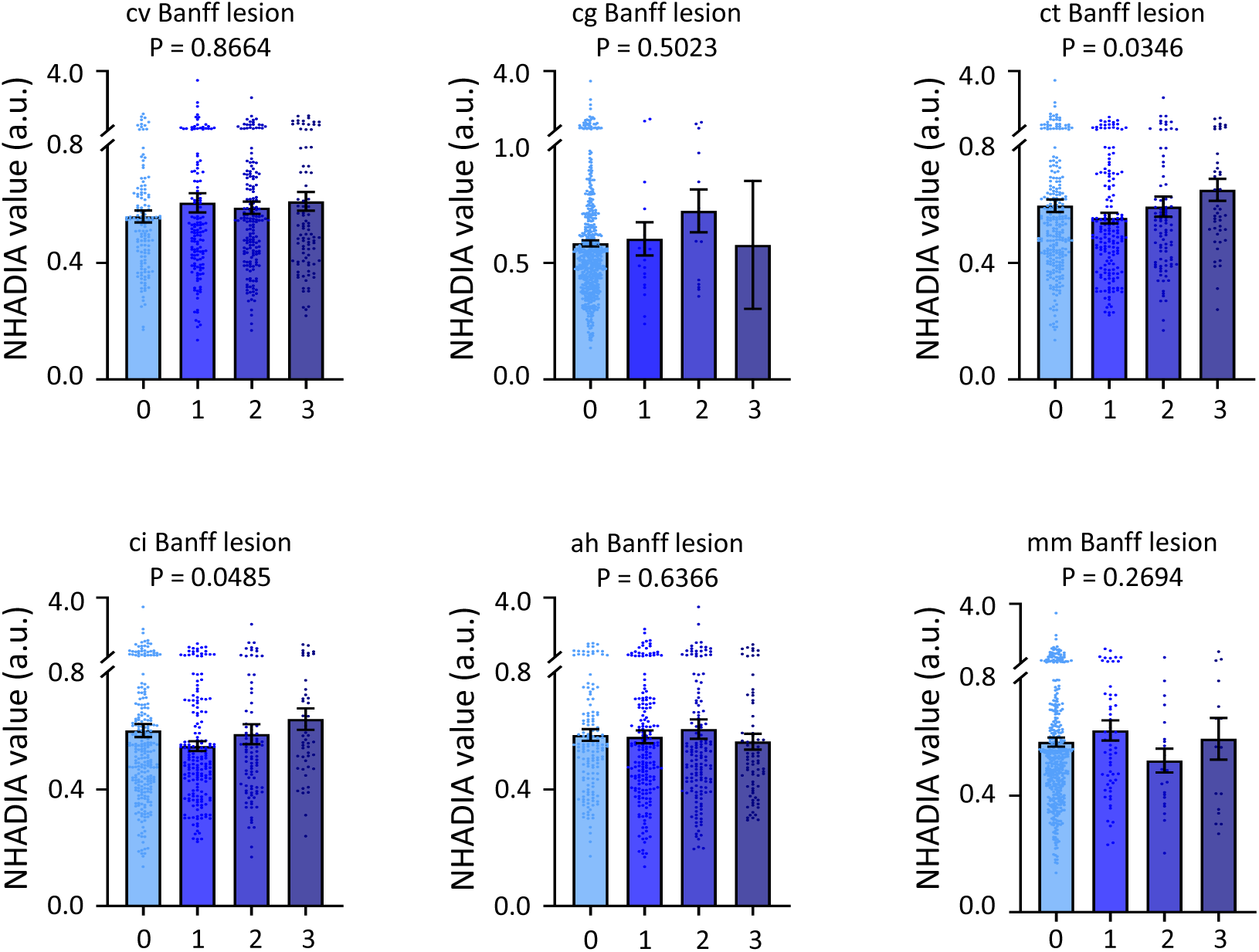
NHADIA values and chronic Banff elementary lesions. Mean NHADIA values (a.u.) according to cv, cg, ct, ci score, ah and mm scores among control and AMRh biopsies. AMRh: histological features of antibody-mediated rejection; NHADIA: Non-HLA Antibody Detection Immunoassay; cv: chronic vascular changes; cg: allograft glomerulopathy; ct: tubular atrophy; ci: interstitial fibrosis; ah: arteriolar hyalinosis; mm: mesangial matrix expansion; P values from two-sided Kruskal-Wallis test. Data are presented as mean values ± SEM.

## Notes

**Funding:** This work was supported by the “Subvention Transplantation et Thérapie Cellulaire 2020 FRM PME20200611626”, the Société Francophone de Transplantation “Bourse de recherche Clinique 2022”, the Société Francophone de Néphrologie, Dialyse et Transplantation, the Day Solvay Foundation and the Boussard Foundation.

### Competing Interest Statement

The authors have declared no competing interest.

