## Supplemental Table 1 for "Non-HLA antibodies worsen the histological phenotype and prognosis of antibody mediated rejection in kidney allografts"

| **Population** | **Total**  **(n = 490)** | **AMRh**  **(n = 326)** | **Control**  **(n = 164)** | ***P* value** |
| --- | --- | --- | --- | --- |
| **Donor characteristics** |  |  |  |  |
| Age, median (IQR) | 55 (44-65) | 57 (45-67) | 50 (41-61) | **0.0013** |
| Sex (Male), n (%) | 245 (50.2) | 162 (50) | 83 (50.6) | 0.9238 |
| Deceased donor, n (%) | 357 (72.9) | 250 (76.7) | 107 (65.2) | **0.0096** |
| ECD (Expanded Criteria Donors), n (%) | 171 (47.9) | 131 (52.4) | 40 (37.4) | **0.0109** |
| Cold ischemia time (h), median (IQR) | 20 (14.33-27.3) | 20 (14.92-27.23) | 19.29 (13.75-27.2) | 0.3261 |
| **Recipient Characteristics** |  |  |  |  |
| Age, median (IQR) | 53 (40-62) | 53 (42-63) | 51 (38-60) | 0.077 |
| Sex (male), n (%) | 297 (60.6) | 199 (61) | 98 (59.8) | 0.8447 |
| Retransplantation, n (%) | 107 (21.8) | 91 (27.9) | 16 (9.8) | **<0.0001** |
| History of pregnancy (female), n (%) | 134 (69.8) | 89 (70.6) | 45 (68.2) | 0.7428 |
| History of blood transfusion, n (%) | 218 (44.5) | 170 (52.1) | 48 (29.3) | **<0.0001** |
| Cause of end stage renal disease |  |  |  |  |
| Glomerulonephritis, n (%) | 113 (23.1) | 72 (22.1) | 41 (25) | 0.4961 |
| Cystic/hereditary/congenital, n (%) | 105 (21.4) | 58 (17.8) | 47 (28.7) | **0.0072** |
| Interstitial nephritis, n (%) | 57 (11.6) | 40 (12.3) | 17 (10.4) | 0.6544 |
| Diabetes, n (%) | 53 (10.8) | 42 (12.9) | 11 (6.7) | **0.0445** |
| Miscellaneous conditions, n (%) | 33 (6.7) | 21 (6.4) | 12 (7.3) | 0.7061 |
| Hypertension, n (%) | 23 (4.7) | 15 (4.6) | 8 (4.9) | 1 |
| Etiology uncertain, n (%) | 106 (21.6) | 78 (23.9) | 28 (17.1) | 0.1032 |
| Delayed graft function, n (%) | 146 (29.9) | 106 (32.5) | 40 (24.5) | 0.1387 |
| **Immunological profile at the index biopsy** |  |  |  |  |
| HLA-A+B+DR mismatches, mean ± SD | 3.3 ± 1.4 | 3.4 ± 1.4 | 3.1 ± 1.4 | 0.0511 |
| HLA-DSAs, n (%) | 292 (59.6) | 219 (67.2) | 73 (44.5) | **<0.0001** |
| **Graft characteristics at time of AMRh** |  |  |  |  |
| Time between transplantation and index biopsy (days), median (IQR) | 344 (67.25-382.75) | 92 (23.25-356.75) | 376 (366-397.25) | **<0.0001** |
| Type of biopsy |  |  |  | **<0.0001** |
| Indication biopsy, n (%) | 181 (36.9) | 181 (55.5) | 0 (0) |  |
| Protocol biopsy, n (%) | 309 (63.1) | 145 (44.5) | 164 (100) |  |
| Serum creatinine at time of biopsy (µmol/L), mean ± SD | 179.6 ± 131.5 | 210 ± 152.5 | 122.4 ± 32.3 | **<0.0001** |
| Proteinuria at time of biopsy (g/g), mean ± SD | 0.5 ± 0.9 | 0.6 ± 1.1 | 0.2 ± 0.5 | **<0.0001** |
