## Supplemental Table 2 for "Non-HLA antibodies worsen the histological phenotype and prognosis of antibody mediated rejection in kidney allografts"

| **Biopsies** |  | **Control**  **(n = 164)** |
| --- | --- | --- |
| *Tubulitis (t)* | Score >0, n (%) | 16 (9.8) |
|  | Score, mean±SD | 0.2 ± 0.6 |
| *Intimal arteritis (v)* |  |  |
|  | Score >0, n (%) | 0 (0) |
|  | Score, mean±SD | 0 ± 0 |
| *Interstitial infiltrate (i)* |  |  |
|  | Score >0, n (%) | 1 (0.6) |
|  | Score, mean±SD | 0 ± 0.1 |
| *Glomerulitis (g)* |  |  |
|  | Score >0, n (%) | 0 (0) |
|  | Score, mean±SD | 0 ± 0 |
| *Peritubular capillaritis (ptc)* |  |  |
|  | Score >0, n (%) | 0 (0) |
|  | Score, mean±SD | 0 ± 0 |
| *Interstitial fibrosis (ci)* |  |  |
|  | Score >0, n (%) | 80 (48.8) |
|  | Score, mean±SD | 0.6 ± 0.8 |
| *Tubular atrophy (ct)* |  |  |
|  | Score >0, n (%) | 75 (46.3) |
|  | Score, mean±SD | 0.6 ± 0.8 |
| *Chronic vascular changes (cv)* |  |  |
|  | Score >0, n (%) | 108 (72) |
|  | Score, mean±SD | 1.2 ± 0.9 |
| *Arteriolar hyalinosis (ah)* |  |  |
|  | Score >0, n (%) | 115 (71) |
|  | Score, mean±SD | 1.1 ± 0.9 |
| *Allograft glomerulopathy (cg)* |  |  |
|  | Score >0, n (%) | 0 (0) |
|  | Score, mean±SD | 0 ± 0 |
| *Mesangial matrix expansion (mm)* |  |  |
|  | Score >0, n (%) | 5 (3.4) |
|  | Score, mean±SD | 0 ± 0.2 |
| *MVI (g+ptc)* |  |  |
|  | Score >0, n (%) | 0 (0) |
|  | Score, mean±SD | 0 ± 0 |
| *C4d* |  |  |
|  | Score >0, n (%) | 0 (0) |
|  | Score, mean±SD | 0 ± 0 |
| *TMA* |  |  |
|  | Yes | 0 (0) |
|  | No | 164 (100) |
