## Supplementary material for "Non-HLA antibodies worsen the histological phenotype and prognosis of antibody mediated rejection in kidney allografts": Table 1

| **Population** | **AMRh**  **(n = 326)** | **HLA-DSA+ AMRh**  **(n = 219)** | **HLA-DSA- AMRh**  **(n = 147)** | ***P* value** |
| --- | --- | --- | --- | --- |
| **Donor characteristics** |  |  |  |  |
| Age, median (IQR) | 57 (45-67) | 55 (44-64) | 59 (49-72) | **0.0053** |
| Sex (Male), n (%) | 162 (50) | 109 (50) | 53 (50) | 1 |
| Deceased donor, n (%) | 250 (76.7) | 169 (77.2) | 81 (75.7) | 0.7814 |
| ECD (Expanded Criteria Donors), n (%) | 131 (52.4) | 83 (49.1) | 48 (59.3) | 0.1393 |
| Cold ischemia time (h), median (IQR) | 20 (14.92-27.23) | 19.83 (14.27-25.72) | 21 (19-28.83) | 0.157 |
| **Recipient Characteristics** |  |  |  |  |
| Age, median (IQR) | 53 (42-63) | 50 (40-61) | 58 (46-66.5) | **0.0004** |
| Sex (male), n (%) | 199 (61) | 132 (60.3) | 67 (62.6) | 0.7178 |
| Retransplantation, n (%) | 91 (27.9) | 80 (36.5) | 11 (10.3) | **< 0.0001** |
| History of pregnancy (female), n (%) | 89 (70.6) | 56 (65.1) | 33 (82.5) | 0.0587 |
| History of blood transfusion, n (%) | 170 (52.1) | 126 (57.5) | 44 (41.1) | **0.0066** |
| Cause of end stage renal disease |  |  |  |  |
| Glomerulonephritis, n (%) | 75 (22) | 50 (22.8) | 22 (20.6 | 0.6725 |
| Cystic/hereditary/congenital, n (%) | 25 (17.8) | 46 (21) | 12 (11.2) | **0.0313** |
| Interstitial nephritis, n (%) | 40 (12.3) | 25 (11.4) | 15 (14) | 0.5899 |
| Diabetes, n (%) | 42 (12.9) | 25 (11.4) | 17 (15.9) | 0.2917 |
| Miscellaneous conditions, n (%) | 21 (6.4) | 12 (5.5) | 9 (8.4) | 0.3404 |
| Hypertension, n (%) | 15 (4.6) | 9 (4.1) | 6 (5.6) | 0.5785 |
| Etiology uncertain, n (%) | 78 (23.9) | 52 (23.7) | 26 (24.3) | 1 |
| Delayed graft function, n (%) | 106 (32.5) | 65 (29.7) | 41 (38.3) | **0.0402** |
| **Immunological profile at the index biopsy** |  |  |  |  |
| HLA-A+B+DR mismatches, mean ± SD | 3.4 ± 1.4 | 3.5 ± 1.4 | 3.3 ± 1.6 | 0.4207 |
| HLA-DSAs, n (%) | 219 (67.2) | 219 (100) | 0 (0) | **< 0.0001** |
| **Graft characteristics at time of AMRh** |  |  |  |  |
| Time between transplantation and index biopsy (days), median (IQR) | 92 (23.25-356.75) | 93 (31.5-373.5) | 85 (17.5-230.5) | **0.0321** |
| Type of biopsy |  |  |  | 0.4766 |
| Indication biopsy, n (%) | 181 (55.5) | 125 (57.1) | 56 (52.3) |  |
| Protocol biopsy, n (%) | 145 (44.5) | 94 (42.9) | 51 (47.7) |  |
| Serum creatinine at time of biopsy (µmol/L), mean ± SD | 210 ± 152.5 | 205.1 ± 147.3 | 220.3 ± 163.3 | 0.889 |
| Proteinuria at time of biopsy (g/g), mean ± SD | 0.6 ± 1.1 | 0.6 ± 1.1 | 0.7 ± 1.1 | 0.5375 |
| **Treatment of graft rejection** |  |  |  |  |
| Anti-thymocyte globulins, n (%) | 13 (4.1) | 6 (2.8) | 7 (6.9) | 0.125 |
| Intravenous corticosteroid, n (%) | 143 (44.8) | 100 (46.1) | 43 (42.2) | 0.5473 |
| IVIG, n (%) | 122 (38.2) | 99 (45.6) | 23 (22.5) | **0.0001** |
| Plasmapheresis, n (%) | 109 (34.2) | 87 (40.1) | 22 (21.6) | **0.001** |
| Rituximab, n (%) | 30 (9.4) | 25 (11.5) | 5 (5) | 0.0663 |
| Eculizumab, n (%) | 11 (3.5) | 9 (4.1) | 2 (2) | 0.5125 |
