## Supplementary material for "Non-HLA antibodies worsen the histological phenotype and prognosis of antibody mediated rejection in kidney allografts": Table 2

| **Biopsies** |  | **AMRh**  **(n = 407)** | **HLA-DSA+ AMRh**  **(n = 290)** | **HLA-DSA- AMRh**  **(n = 117)** | ***P* value** |
| --- | --- | --- | --- | --- | --- |
| *Tubulitis (t)* | Score >0, n (%) | 127 (31.3%) | 86 (29.8%) | 41 (35%) | 0.3445 |
|  | Score, mean±SD | 0.6+-1 | 0.5+-0.9 | 0.8+-1.2 | 0.1583 |
| *Intimal arteritis (v)* |  |  |  |  |  |
|  | Score >0, n (%) | 43 (10.6%) | 23 (7.9%) | 20 (17.1%) | **0.0116** |
|  | Score, mean±SD | 0.2+-0.5 | 0.1+-0.5 | 0.3+-0.6 | **0.0064** |
| *Interstitial infiltrate (i)* |  |  |  |  |  |
|  | Score >0, n (%) | 55 (13.5%) | 33 (11.4%) | 22 (18.8%) | 0.0554 |
|  | Score, mean±SD | 0.2+-0.5 | 0.1+-0.5 | 0.3+-0.7 | **0.0393** |
| *Glomerulitis (g)* |  |  |  |  |  |
|  | Score >0, n (%) | 330 (81.1%) | 238 (82.1%) | 92 (78.6%) | 0.4845 |
|  | Score, mean±SD | 1.3+-0.9 | 1.4+-0.9 | 1.1+-0.8 | **0.0112** |
| *Peritubular capillaritis (ptc)* |  |  |  |  |  |
|  | Score >0, n (%) | 302 (74.2%) | 224 (77.2%) | 78 (66.7%) | **0.0332** |
|  | Score, mean±SD | 1.4+-1 | 1.5+-1 | 1.2+-1 | **0.003** |
| *Interstitial fibrosis (ci)* |  |  |  |  |  |
|  | Score >0, n (%) | 239 (58.7%) | 172 (59.3%) | 67 (57.3%) | 0.7391 |
|  | Score, mean±SD | 1+-1 | 1+-1 | 1+-1 | 0.9722 |
| *Tubular atrophy (ct)* |  |  |  |  |  |
|  | Score >0, n (%) | 232 (57.1%) | 166 (57.4%) | 66 (56.4%) | 0.9119 |
|  | Score, mean±SD | 1+-1 | 1+-1 | 1+-1 | 0.9736 |
| *Chronic vascular changes (cv)* |  |  |  |  |  |
|  | Score >0, n (%) | 300 (76.3%) | 204 (73.4%) | 96 (83.5%) | **0.0367** |
|  | Score, mean±SD | 1.5+-1.1 | 1.4+-1.1 | 1.7+-1 | **0.0045** |
| *Arteriolar hyalinosis (ah)* |  |  |  |  |  |
|  | Score >0, n (%) | 301 (75.2%) | 211 (74.6%) | 90 (76.9%) | 0.7027 |
|  | Score, mean±SD | 1.4+-1 | 1.3+-1 | 1.4+-1.1 | 0.3947 |
| *Allograft glomerulopathy (cg)* |  |  |  |  |  |
|  | Score >0, n (%) | 36 (9%) | 30 (10.5%) | 6 (5.3%) | 0.1218 |
|  | Score, mean±SD | 0.1+-0.5 | 0.2+-0.5 | 0.1+-0.4 | 0.1119 |
| *Mesangial matrix expansion (mm)* |  |  |  |  |  |
|  | Score >0, n (%) | 94 (26.1%) | 67 (26.5%) | 27 (25.2%) | 0.8957 |
|  | Score, mean±SD | 0.4+-0.8 | 0.4+-0.8 | 0.4+-0.8 | 0.9015 |
| *MVI (g+ptc)* |  |  |  |  |  |
|  | Score >0, n (%) | 398 (97.8%) | 286 (98.6%) | 112 (95.7%) | 0.1275 |
|  | Score, mean±SD | 2.7+-1.3 | 2.9+-1.3 | 2.3+-1.3 | **0.0001** |
| *C4d* |  |  |  |  |  |
|  | Score >0, n (%) | 197 (48.4%) | 134 (46.2%) | 63 (53.8%) | 0.1886 |
|  | Score, mean±SD | 1.1+-1.3 | 1.1+-1.3 | 1.2+-1.2 | 0.2372 |
| *C4d detail* |  |  |  |  |  |
|  | 0 | 210 (51.6%) | 156 (53.8%) | 54 (46.2%) | 0.1886 |
|  | 1 | 31 (7.6%) | 23 (7.9%) | 8 (6.8%) | 0.8375 |
|  | 2 | 76 (18.7%) | 47 (16.2%) | 29 (24.8%) | **0.0497** |
|  | 3 | 90 (22.1%) | 64 (22.1%) | 26 (22.2%) | 1 |
| *TMA* |  |  |  |  | 0.524 |
|  | Yes | 29 (7.1%) | 19 (6.6%) | 10 (8.5%) |  |
|  | No | 378 (92.9%) | 271 (93.4%) | 107 (91.5%) |  |
