## Supplemental M&M for "Non-HLA antibodies worsen the histological phenotype and prognosis of antibody mediated rejection in kidney allografts"

**Supplementary Methods**

***Detection of HLA-DSA***

All patients were tested for the presence of HLA-DSA at 3 months, at 12 months and for clinical indications. The presence of circulating HLA-DSA against HLA-A, HLA-B, HLA-Cw, HLA-DR, HLA-DQ, and HLA-DP was determined using single-antigen flow bead assays (One Lambda, Inc., Canoga Park, CA) on a Luminex platform. Beads with a normalized mean fluorescence intensity (MFI) greater than 500 arbitrary units were considered positive.

***NHADIA***

Serum samples collected at the time of biopsies after transplantation were tested with the NHADIA. After washing with PBS, differentiated CiGEnCΔHLA cells were trypsinized (TrypLE Express, Thermo Fisher) and washed before incubation with a fixable viability dye (Thermo Fisher) for 25 minutes at 4°C. Then, the CiGEnCΔHLA cells were incubated with patient sera diluted 1:2 in PBS containing 0.05% BSA and 2 mM EDTA for 30 minutes at 4°C. For the negative control, cells were incubated with PBS only. For the positive control, cells were incubated with a serum containing 33 known non-HLA Abs (LABScreen Autoantibody LSAUT-PC, One Lambda). After two more washes, the cells were incubated with an Alexa Fluor® 488-conjugated anti-human IgG Ab (AffiniPure F(ab')₂ Fragment Donkey Anti-Human IgG (H+L), Interchim) for 30 minutes at 4°C. Finally, the cells were fixed with PBS containing 4% Formaldehyde. Fluorescence was measured by flow cytometry (LSR Fortessa X-20, BD Biosciences), and geometric means of fluorescence intensity (Geo MFI) were calculated using Kaluza software v2.1 (Beckman Coulter). The results were calculated as the ratio of the Geo MFI_sample_ to the Geo MFI_positivecontrol_. This Non-HLA Antibody Immuno Assay (NHADIA) using the CiGEnCΔHLA cells (**Figure 2B**) was patented (patent n°EP21 305 960.3 (BIO21230)).
